# New detection of SARS-CoV-2 in two cats height months after COVID-19 outbreak appearance in France

**DOI:** 10.1101/2021.03.24.436830

**Authors:** Matthieu Fritz, Nicolas Nesi, Solène Denolly, Bertrand Boson, Vincent Legros, Serge G. Rosolen, Alexandra Briend-Marchal, Meriadeg Ar Gouilh, Eric M. Leroy

## Abstract

Although there are several reports in the literature of SARS-CoV-2 infection in cats, few SARS-CoV-2 sequences from infected cats have been published. In this report, SARS-CoV-2 infection was evaluated in two cats by clinical observation, molecular biology (qPCR and NGS), and serology (Microsphere immunoassay and seroneutralization). Following the observation of symptomatic SARS-CoV-2-infection in two cats, infection status was confirmed by RT-qPCR and, in one cat, serological analysis for antibodies against N-protein and S-protein, as well as neutralizing antibodies. Comparative analysis of five SARS-CoV-2 sequence-fragments obtained from one of the cats showed that this infection was not with one of the three recently emerged variants of SARS-CoV-2. This study provides additional information on the clinical, molecular, and serological aspects of SARS-CoV-2 infection in cats.

## Introduction

SARS-CoV-2 has shown relatively generalist capacities by infecting many animal species, making it a good-model in One Health research (MacLean et al., 2021). Indeed, SARS-CoV-2 infections have been detected in numerous animal species living in close contact with infected humans. Based on RNA detection, serological studies, and experimental infections, numerous animal species have proven susceptibility to SARS-CoV-2 (Shi et al., 2020), including *Mustelidae* (ferret, mink) (Oude Munnink et al., 2020), *Canidae* (dog) (Sit et al., 2020), *Felidae* (cat, tiger, lion) (Sailleau et al., 2020) and *Cricetidae* (hamster, rat, mouse) (Sia et al., 2020) as well as by the highly transmissible British variant (B.1.1.7) (Ferasin et al., 2021). On March 24^th^, the World Organisation for Animal Health reports cases of SARS-CoV-2 infections in cats in 17 countries (USA, China, Belgium, Germany, Spain, France, Russia, United-Kingdom, Japan, Italy, Chile, Brazil, Greece, Canada, Argentina, Switzerland, and Latvia). In the medical and scientific literature, we found 23 papers examining natural SARS-CoV-2 infection in a total of 2242 cats (Barrs et al., 2020; Carlos et al., 2021; Chen, Huang, Zhang, Zhang, & Jin, 2020; Deng et al., 2020; Ferasin et al., 2021; Fritz et al., 2021; Garigliany et al., 2020; Hamer et al., 2020; Hosie et al., 2020; Klaus et al., 2021; Michelitsch, Hoffmann, Wernike, & Beer, 2020; Musso et al., 2020; Neira et al., 2020; Newman et al., 2020; Pagani et al., 2021; Patterson et al., 2020; Ruiz-Arrondo et al., 2020; Sailleau et al., 2020; Segalés et al., 2020; Stevanovic et al., 2020; Temmam et al., 2020; Villanueva-Saz et al., 2021; Zhang et al., 2020). Among these cats, only 94 were positive for ongoing or previous SARS-CoV-2 infection, as detected by qPCR or by serology. Additionally, the U.S Department of Agriculture has reported 67 cases of SARS-CoV-2 infection in cats identified by at least one of these assays. Viral RNA detected in 25 cats suggested that COVID-19 infections were transmitted from infected owners. These few sequences are available (31 on Global Initiative on Sharing All Influenza Data (Gisaid) and 18 on Genbank) (https://www.gisaid.org/; https://www.ncbi.nlm.nih.gov/genbank/). While most cases of SARS-CoV-2 infection in cats were asymptomatic, some cats (14/94) experienced lethargy, mild respiratory or digestive symptoms (sneezing, coughing, ocular discharge, vomiting, and anorexia), and two studies reported severe respiratory problems in two cats (Garigliany et al., 2020; Musso et al., 2020). Recently, a study has shown association between B.1.1.7 infection and clinical signs of myocarditis in cats (Ferasin et al., 2021). Moreover, several experimental studies have shown that SARS-CoV-2 can be transmitted between cats (Bosco-Lauth et al., 2020; Gaudreault et al., 2020; Halfmann et al., 2020). As yet, there is no evidence of cat-to-human transmission. Here we present a clinical and biological investigation of SARS-CoV-2 infection of two cats that presented with mildly symptomatic disease, and that came from households with confirmed cases of COVID-19 sampled during the second wave (October-November 2020) of infections in France.

## Materials and methods

### First line diagnostic: Veterinary diagnostic laboratory (VEBIO)

#### RNA extraction

RNA extraction from nasopharyngeal and rectal swabs was done using QIAamp Viral RNA Mini Kit (QIAGEN). Swabs were resuspended in 200μl of ATL + 20μl of proteinase K then heated at 70°C for 10 min. 200μl of ATL + 20μl of proteinase K was then added, and again heated to 70°C. Finally, 200μl of ethanol was added and the entire volume (640μl) transferred to a column. Subsequent steps proceeded according to the manufacturer’s protocol.

#### Real-time rever se-transcription PCR

Viral RNA was quantified using real the Genesig® Real Time PCR Coronavirus COVID-19 (CE IVD) kit (Primer Design) using the manufacturer’s protocol and QuantStudio 5 Real-Time PCR System thermocycler (Thermo Fisher Scientific)

### Second-line diagnostic: Virology laboratory of Caen Hospital

#### RNA extraction and Removal of genomic DNA

Nasopharyngeal and rectal swabs were resuspended in 500μl of PBS, of which 140μl was used for RNA extraction using the EZ1 RNA Tissue Mini kit (QIAGEN). The homogenate was then treated with the Turbo DNA-free kit (Thermo Fisher Scientific), according to the manufacturer’s protocol, in order to remove genomic DNA.

#### Ribosomal RNA Depletion and Real-time reverse-transcription PCR

Critical to enable cost-effective sequencing of RNA samples, we depleted the eukaryotic rRNA using the NEBNext rRNA Depletion Kit (Human/Mouse/Rat). After rRNA depletion, double-stranded cDNA was synthesized by real-time reverse-transcription (RT-PCR) using Superscript III platinium One-step Quantitative RT-PCR System (Invitrogen) as described previously with minor modifications (Corman et al., 2020). Briefly, a 25 μL reaction contained 5 μL of RNA, 12.5 μL of 2 × reaction buffer, 1 μL of reverse transcriptase/ Taq mixture, 0.4 μL of a 50 mM magnesium sulphate solution (Invitrogen), 1μl of a 10μM primer, and 0.5l μl of a 10μM probe. Thermal cycling was performed at 50 °C for 15 min for reverse transcription, followed by 95 °C for 2 min and then 45 cycles of 95 °C for 15 s and 60 °C for 30 s using a Light Cycler 480 (Roche).

To increase the amount of genetic material available for sequencing, whole transcriptome amplification (WTA) was performed with QuantiTect Whole Transcriptome kit (QIAGEN) according to the manufacturer’s instructions. Amplified DNA reactions was then purified using AMPure XP beads and quantified using Qbit dsDNA BR Assay Kit on a Qubit 3.0 fluorimeter (Invitrogen).

In complement to the *de novo* approach, we also used an adapted version of the published protocol from the ARTIC Network using ARTIC (Quick, 2020) primer scheme version 3 which produces ∼400 bp overlapping amplicons over the SARS-CoV-2 genome.

#### ONT library preparation and MinION sequencing

Libraries were prepared without shearing to maximize sequencing read length. The Oxford Nanopore Technology (ONT) protocol for native barcoding genomic DNA sequencing was followed using the barcoded ligation sequencing kit SQK-LSK108 and the EXP-NBD104 Native Barcoding kit. Sequencing libraries were constructed, and sequencing performed according to manufacturer’s instructions, as briefly described below. First, the NEBNext Ultra II End Repair/dATailing module (E7546S, NEB, USA) was used to prepare 1000 ng DNA samples. End-prepared DNA was ligated with native barcode adapters NBD04 using Blunt/TA Ligase Master Mix (M0367S, NEB, USA). Following the barcode ligation reaction, the DNA was cleaned with AMPure XP beads. The two samples were then pooled to produce a 54 μl equimass pool used for adapter ligation with the ‘Native Barcoding Adapter Mix (BAM). The final library was loaded onto an R9.4 flowcell (FLO-MIN106, Oxford Nanopore Technologies, UK), and the run was performed on a MinION Mk1B device (ONT).

#### Genome assembly

Following the MinION run, reads generated were basecalled and subsequently demultiplexed using Guppy GPU basecaller and barecoder (Oxford Nanopore Technologies). Reads were then mapped against a custom reference of SARS-CoV-2 genome comprising four Chinese and 70 early French sequences using Bowtie2 (Langmead & Salzberg, 2012) and minimap2 (Heng Li, 2018). Finally, a consensus genome sequence based on mapped reads was generated with bcftools consensus (H. Li, 2011). SARS-CoV-2 sequences were deposited on GISAID (EPI_ISL_1328819;EPI_ISL_1328821; EPI_ISL_1328824; EPI_ISL_1328826).

#### Microsphere immunoassay (MIA)

MIA was proceeded in Maladies Infectieuses et vecteurs: Ecologie, Génétique, Evolution et Contrôle (MIVEGEC). Cat serum samples were tested units using a multiplex Microsphere immunoassay (MIA). 10μg of three recombinant SARS-CoV-2 antigens: nucleoprotein (N), receptor-binding domain (RBD), and trimeric spike (tri-S) were used to capture specific serum antibodies whereas a human protein (O6-methylguanine DNA methyltransferase) was used as a control antigen in the assay. Distinct MagPlex microsphere sets (Luminex Corp) were respectively coupled to viral antigens using the amine coupling kit (Bio-Rad Laboratories) according to manufacturer’s instructions. The MIA procedure was performed as described previously (Fritz et al., 2021). Briefly, microsphere mixtures were successively incubated with serum samples (1:400) biotinylated protein A and biotinylated protein G (4 μg/ml each) (Thermo Fisher Scientific), and Streptavidin-R-Phycoerythrin (4 μg/ml) (Life technologies) on an orbital shaker and protected from the light. Measurements were performed using a Magpix instrument (Luminex). To account for nonspecific binding of antibodies to beads, Relative Fluorescence Intensities (RFI) were calculated for each sample by dividing the Mean Fluorescence Intensity (MFI) signal measured for the antigen-coated microspheres by the MFI signal obtained for the control microspheres. Specific seropositivity cut-off values for each antigen were set at three standard deviations above the mean RFI of the 18 dogs and 14 cat serum samples sampled before 2019. Based on a pre-pandemic population, MIA specificity was set at 100% for dogs and cats.

#### Neutralization activity measurement

Neutralization activity measurement was proceeded in Virus enveloppés, vecteurs et immunothérapie (EVIR) team from CIRI. An MLV-based pseudoparticle carrying a GFP reporter pseudotyped with SARS-CoV-2 spike protein (SARS-CoV-2pp) was used to measure the neutralizing antibody activity in cat sera. Each SARS-CoV-2 positive sample detected by MIA was processed according to a neutralization procedure as previously described (Legros et al., 2021). The level of infectivity is expressed as the percentage of GFP-positive cells and compared to cells infected with SARS-CoV-2pp incubated without serum. Pre-pandemic sera from France was used as negative controls, and anti-SARS-CoV-2 RBD antibody was used as a positive control.

## Results

Cat 1 was a solitary and sedentary 5-year-old female European whose only contact was with her owner. Her last vaccination was 3 years prior and she was presented with no previous medical history. On October 24^th^ 2020, 10 days after her owner developed symptoms and tested positive for SARS-CoV-2 infection, Cat 1 developed sneezing with non-purulent nasal secretions but no digestive trouble or others notable symptoms. Five days later, on October 29, clinical examination of the cat showed pink mucous membranes, a heart’s rate of 175 bpm (110bpm-200bpm), a rectal temperature of 38.4°C (38°C-39°C), slight dehydration, a urea seric concentration of 0.49g/l (0.4g/l-0.6g/l), and a creatinine seric concentration of 12.7mg/l (10mg/l-20mg/l). A 10-days treatment of 1 Doxybactin tablet in the evening and a daily dose of Meloxoral taken over 3 days with the morning meal was then initiated. An improvement of the symptoms was observed in this cat three days after the onset of the treatment. Based on the owner’s positive SARS-CoV-2 diagnosis and the cat’s symptoms, a blood sample and two oropharyngeal and rectal swabs were collected for further SARS-CoV-2 testing. Swabs were submitted to VEBIO for RNA SARS-CoV-2 detection. The oropharyngeal swab tested positive by RT-qPCR targeting the ORF1ab with a ct value of 20.41. No viral RNA was detected in the rectal swabs. The virology laboratory of Caen Hospital confirmed the diagnosis following detection of viral RNA in the second ororpharyngeal swab by RT-qPCR targeting gene E with a ct value of 21.43 (Table 1). We did not obtain a SARS-CoV-2 sequence due to the sample’s poor conservation condition prior to its arrival at the lab. To detect anti-SARS-CoV-2 IgG antibodies, the cat 1 serum was analysed by the MIVEGEC and EVIR teams, using MIA and retrovirus-based pseudoparticle assay. Antibodies against N, RBD, and tri-S SARS-CoV-2 proteins as well as neutralizing antibodies were detected, confirming productive infection in cat 1 (Table 1).

**Table 1.** Characteristics of cats sampled during the second wave of SARS-CoV2 infections in France

| Cat ID | Age | Gender | Severity of cat's symptoms | Nasopharyngeal swab | Rectal swab | Serology (MIA) | Serology (Seroneutralisation) |
| --- | --- | --- | --- | --- | --- | --- | --- |
| Cat 1 | 5 | F | Mild | Positive | Negative | Positive | Positive |
| Cat 2 | 13 | M | Mild | Positive | Negative | ND | ND |

Cat 2 was a 13-year-old male European with chronic rhinitis and living with two other cats. The cat’s owner, who had recently tested positive for SARS-CoV-2, reported an acute deterioration in his three cats’ general condition, without others details. A November 20th, 2020 clinical investigation reported retro-mandibular adenopathy, and no other symptoms was observed. As with Cat 1, oropharyngeal and rectal swabs were collected but with no blood sample. Similarly, VEBIO and Caen laboratories detected viral RNA in both oropharyngeal swabs, with ct-values of 20.55 and 23.44, respectively (Table 1). No viral RNA was detected in the rectal swabs. Again due to poor conservation of the swabs prior to their arrival in our laboratory, only 5 partial fragments of the SARS-CoV-2 genome could be obtained by high-throughput sequencing on RNA derived from the oropharyngeal swabs. In fragment 2, we did not observe the 11288-11296 deletion characterizing 3 variants of concern (English B1.1.7, South African B.1.351, and Brazilian P.1) (Table 2), indicating that this cat was not infected by any of these three novel variants. Compared to the original Wuhan_Hu-1 reference sequence, we observed, in fragment 4, only one single nucleotide polymorphism (SNP) -G25563T-leading to the amino acid change Q57H. This H57 mutation is widely distributed around the world and was present in about 70% of the sequences reported in France between October and December 2020, according to Nextstrain (https://nextstrain.org/).

**Table 2:**
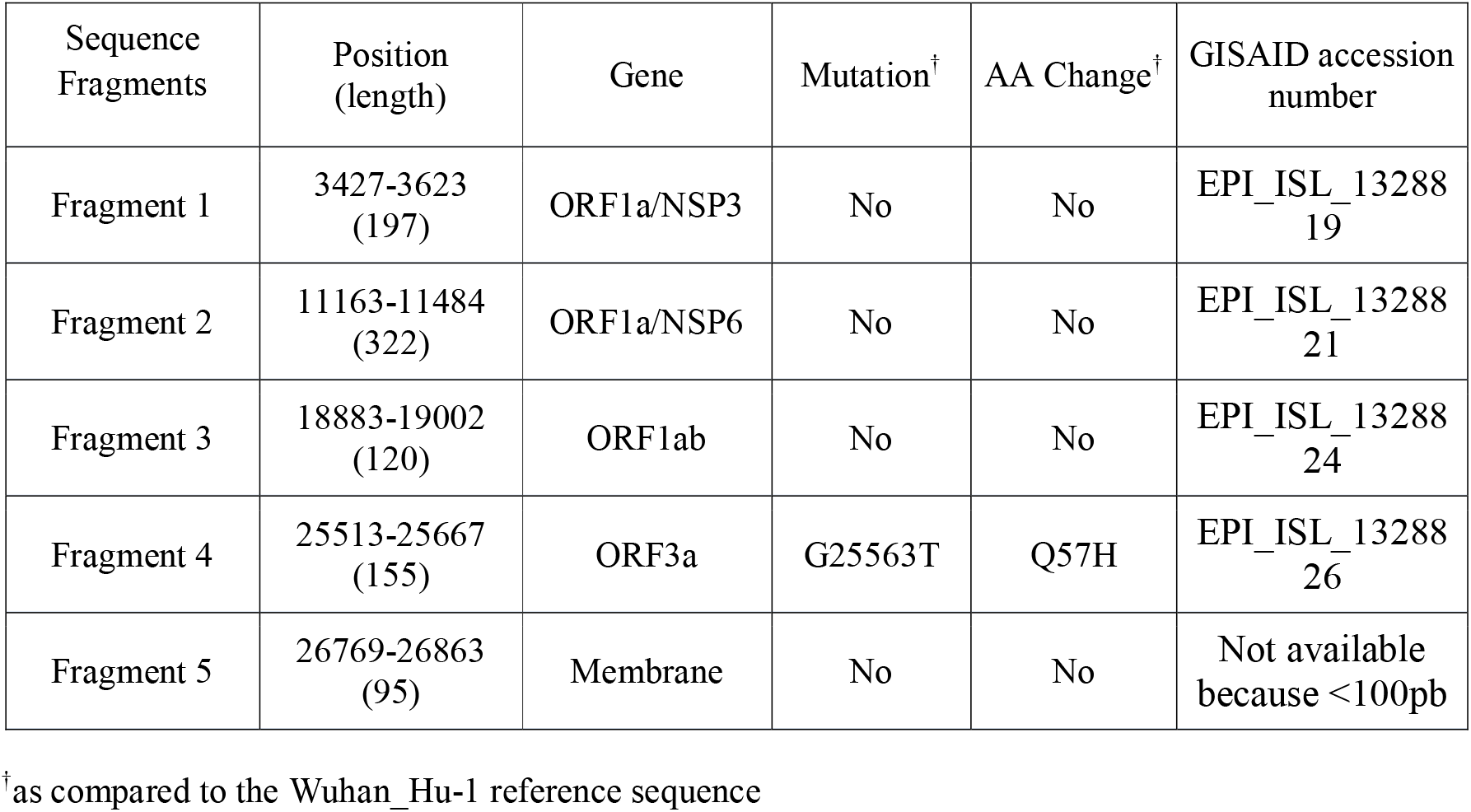
Details of the five SARS-CoV2 genome fragments obtained from cat 2

## Discussion

Here we report SARS-CoV-2 infection in two cats, diagnosed by molecular and serological assays, and with mild clinical manifestations, including sneezing for one cat. Although we cannot definitively rule out infection of the cats by an individual outside the household, the information given by COVID19+ owners, including the exclusive and unique contact with its owner for cat 1 and the general deterioration in the condition of all cats of the cat 2 owner, strongly suggests a transmission from owners to cats.

Human-to-cat transmission is now widely reported worldwide, and while certainly not insignificant in COVID19+ households, precise estimates of the frequency of such transmission has not been established (Fritz et al., 2021; Hamer et al., 2020). To our knowledge, among the 23 papers investigating possible SARS-CoV-2 infection in cats, only 14 reported PCR results confirming infection in 27 cats. The present study is only the second showing molecular results of cat infection in France, eight months after the first case report by Saillau *et al*. (Sailleau et al., 2020). It appears that the majority of infections in cats are asymptomatic, but can, at times, lead to lethargy, mild respiratory or digestive disease and, rarely, acute respiratory clinical signs (as observed in two cats). We report here one case of mild respiratory symptoms (sneezing) in a cat, an observation consistent with symptomatic infections sporadically observed in other studies. We did not observe any digestive symptoms, as previously reported in a French cat. In accordance with other recent studies, our results suggest that since the beginning of SARS-CoV-2 pandemic, almost one year after the first report of infection in a cat, the pathology in cats has not changed globally, with only a relatively small proportion of cases reported as a result of clinical investigation by a veterinarian. This low pathogenicity can explain the paucity of studies reporting SARS-CoV-2 infection in cats in the absence of a global pet detection policy.

The two cats in the study were sampled during the second wave of infection in France and at the beginning of the ongoing emergence of multiple novel variants. Although we did not find evidence of infection by one of the three novel variants in cat 2, the emergence of these new variants raises the question of potential changes in pathogenicity or transmissibility in domestic animals. This question will become rapidly crucial in a very near future as the British variant, known to be much more infectious, is currently removing the ancestral variant of SARS-CoV-2 in France as well in other countries of Europe. Therefore, it is becoming more and more important to implement a One Health approach to face SARS-CoV-2 epidemic that takes into account infection and viral circulation in pets.

## Acknowledgements

We are grateful to the pet owners for giving us their permission to sample their pets. We thank veterinarians that helped us with sampling. We thank Estelle Leperchois, Simon Thierry and Trung Thanh Nguyen for technical support and assistance. We are grateful to François-Loïc Cosset for helpful discussions and critical reading of this paper. We also thank Kurt McKean for English editing of the manuscript (https://octopusediting.com/).

## Data Availability Statement

The data that support the findings of this study are available from the corresponding author upon reasonable request.

